# Robust Identification of Temporal Biomarkers in Longitudinal Omics Studies

**DOI:** 10.1101/2021.11.19.469350

**Authors:** Ahmed A. Metwally, Tom Zhang, Si Wu, Ryan Kellogg, Wenyu Zhou, Hua Tang, Michael Snyder

## Abstract

Longitudinal studies increasingly collect rich ‘omics’ data sampled frequently over time and across large cohorts to capture dynamic health fluctuations and disease transitions. However, the generation of longitudinal omics data has preceded the development of analysis tools that can efficiently extract insights from such data. In particular, there is a need for statistical frameworks that can identify not only which omics features are differentially regulated between groups but also over what time intervals. Additionally, longitudinal omics data may have inconsistencies, including nonuniform sampling intervals, missing data points, subject dropout, and differing numbers of samples per subject. In this work, we developed a statistical method that provides robust identification of time intervals of temporal omics biomarkers. The proposed method is based on a semi-parametric approach, in which we use smoothing splines to model longitudinal data and infer significant time intervals of omics features based on an empirical distribution constructed through a permutation procedure. We benchmarked the proposed method on five simulated datasets with diverse temporal patterns, and the method showed specificity greater than 0.99 and sensitivity greater than 0.72. Applying the proposed method to the Integrative Personal Omics Profiling (iPOP) cohort revealed temporal patterns of amino acids, lipids, and hormone metabolites that are differentially regulated in male versus female subjects following a respiratory infection. In addition, we applied the longitudinal multi-omics dataset of pregnant women with and without preeclampsia, and the method identified potential lipid markers that are temporally significantly different between the two groups. We provide an open-source R package, *OmicsLonDA* (Omics Longitudinal Differential Analysis): https://bioconductor.org/packages/OmicsLonDA to enable widespread use.

## 1. Introduction

Human health is highly dynamic, and there is great interest in better understanding how wellness and disease states fluctuate over time in relation to different variables such as lifestyle or treatment perturbations. While genomics provides a blueprint for life, health states are also reflected by many other ‘omics’ such as transcriptomics, proteomics, metabolomics, lipidomics, microbiomics. With rapid advances and decreasing costs in sequencing and mass spectrometry, many studies are beginning to measure comprehensive omics profiles at frequent timepoints across many individuals. Longitudinal omics studies generate enormous datasets; however, there is currently a major bottleneck in analyzing this data to extract and interpret meaningful findings. In particular, there is a need for robust statistical methods for longitudinal omics.

Longitudinal omics data has its own properties that differentiate it from cross-sectional experiments, including high dimensional feature space, temporal and intrapersonal variation, and samples characterized by heterogeneity of various natures. These heterogeneities include a different number of samples per subject, uncaptured data points, variable time of sample collection “sampled nonuniformly”, and omics features often represent a biological process that usually exhibits temporal variation. Another aspect is the variability in temporal dependence structure, “variance-covariance structure”, between repeated measurements. All of these characteristics of longitudinal omics data make the analysis a challenging task. Methods developed for longitudinal omics data analysis can be categorized into the following groups: (a) Methods that extract omics biomarkers for a specific phenotype ^1^, (b) Methods that build mechanistic models to describe the underlying mechanism involved in gene regulation, metabolism, or protein-protein-interaction causally related to specific phenotype ^2–4^, (c) Identifying clusters of omics features that have similar expression patterns ^5^.

For the class of methods that identify omics biomarkers, many statistical models have been proposed. The joint mixed model, which is widely used, links separate linear mixed models by allowing their model-specific random effects to be correlated ^6^. The advantages of this approach include well-established theory and efficiency gains ^7,8^. More importantly, a joint random-effect model allows the correlation between different outcomes to be assessed. It can provide a succinct summary of not only how the evolution of one outcome variable is correlated to the evolution of another outcome, but also how the correlation between outcomes changes over time (‘evolution of association’) ^9^. On the other hand, the mixed-effect model comes with its set of assumptions, such as homogeneity of variance of the residuals being equal across groups ^10^ and normality of the residuals ^11^. In many situations, these assumptions are violated. With the rapidly increasing size and complexity of omics datasets, nonparametric methods ^12,13^ are emerging as the primary methods for biomedical analysis. Nonparametric statistics have the advantage of making minimal distributional assumptions and can scale to fit the complexity of the data. A recent non-parametric robust method, *bootLong*, was developed for extracting microbial biomarkers from longitudinal microbiome data based on a moving block bootstrap approach ^14^. It accounts for within-subject dependency by using overlapping blocks of repeated observations within each subject. It then infers biomarkers based on approximately pivotal statistics. Although *bootLong* shows promising results in identifying microbial biomarkers in microbial longitudinal studies, it does not provide time intervals of differences between the study phenotypes. Another method, *MetaLonDA*, has been proposed to find time intervals of significant microbial biomarkers using a permutation test ^15^. *MetaLonDA* is tailored to microbial experiments through the use of a negative binomial distribution.

In this paper, we introduce a robust method to perform longitudinal differential analysis on omics features in order to identify time intervals of differences between study groups. The method is based on a semi-parametric approach, where we use smoothing splines to model longitudinal data and infer significant time intervals of changes in omics features based on an empirical distribution constructed through a permutation procedure. The proposed method can handle all types of inconsistencies in sample collections and adjust for subjects’ specific baseline. Identifying biomarkers and their significant time interval differences can inform intervention strategies (drugs, probiotics, antibiotics, supplements), and most importantly, may indicate the best time for interventions to be administered to patients. The method achieved a correctly calibrated type-I error rate and is robust to data collection inconsistencies that commonly occur in longitudinal human studies. Application of the proposed method to iPOP cohort revealed a multitude of sex differences in dynamic respiratory infection response. To our knowledge this is the first study to investigate sexual dimorphism in infection response with frequent temporal sampling and delineation of the dynamic infection response for each sex. We also applied *OmicsLonDA* on a longitudinal lipidomics study on preeclampsia for the identification of time intervals that lipids are significantly different between pregnancy with and without preeclampsia. We provide an open-source Bioconductor R package, *OmicsLonDA* (Omics Longitudinal Differential Analysis), for widespread availability.

## 2. Methods

The proposed method aims to find the feature’s significant time intervals (FSTI) of differences between each pair of the tested groups (e.g., healthy vs. diseased, male vs. female, etc.). The method works on unpaired experiment design, where subjects’ longitudinal samples are related to only one of the tested groups. We model the longitudinal data in a time series model using a spline kernel. Although, in theory, longitudinal data should be correlated, the 1st order auto-correlation is not that high. This is mainly due to the fact that longitudinal samples are taken far apart from each other. For this reason, we do not consider autocorrelation in our model due to the complexity of assuming a valid dependency structure. The input data to the method is the processed (filtered based on quality control thresholds, annotated, quantified, normalized, corrected for batch effect and sequencing depth) measurements of any of the omics experiments, such as genes expression from RNAseq experiments ^16^, proteins levels from proteomics experiments ^17^, metabolites intensities from metabolomics experiments ^18^, microbial abundance from metagenomics experiments ^19^. The data processing output of each of these omics assays can be summarized in a matrix *C* with a dimension of *m*×*n* where *m* denotes the number of omic features and *n* denotes the number of samples. *C(i,j)* represents the quantity from sample *j* that is annotated to feature *i*. The proposed method is based on 4 main steps as shown in **Figure 1**: (a) adjust measurements based on each subject’s profile, (b) fitting Gaussian smoothing spline regression model, (c) permutation test to generate an empirical distribution of the test statistic of each time interval, and (d) inference of significant time intervals of omics features. The details of the method are described in the following sections.

**Figure 1:**
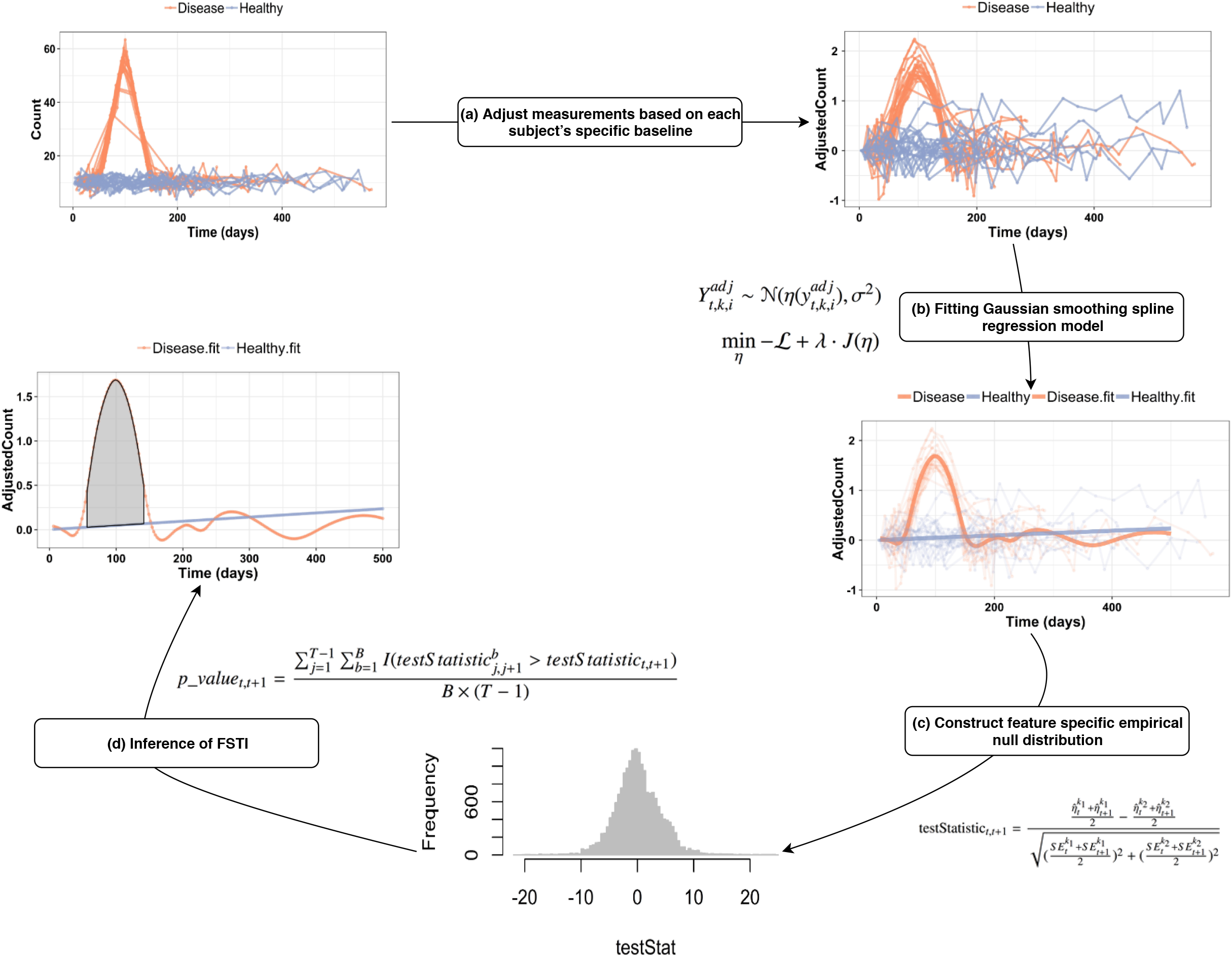
Overview of the main steps of the proposed method: adjust measurements based on each subject’s specific baseline, global testing using to select candidate features for time intervals analysis, fitting Gaussian smoothing spline regression model, permutation test to generate an empirical distribution of the test statistic, and inference of feature’s significant time intervals (FSTI).

### 2.1 Adjusting for subject’s personal profile

Interpersonal omics values can vary dramatically between subjects. Usually, people cluster according to themselves ^20^. Hence, there is a need to adjust longitudinal samples based on the subject’s profile. In this work, we implemented two techniques for adjusting personal profiles. The first strategy is based on using the first sample of the study as the baseline and adjusting each following sample to the baseline. The baseline timepoint is usually chosen to be the sample prior to perturbation (e.g., infection, surgery, infection, etc.), or at a steady-state condition. This strategy is effective when the baseline timepoint is right before the perturbations. For each omic feature *f* under consideration, we first adjust for the difference in the personal baseline. Our strategy is to calculate the log-ratio between omic feature’s level of each timepoint *t* to the level of the same omic feature at the subject’s chosen baseline *tb* (**Eq.1**), where *y*_*i,t*_ is the measure of the omic feature of subject *i* at time point *t*, and *t*_*b*_ is the *ith* subject’s baseline. Besides adjusting for the personal baseline, the logged ratio reduces the positive skewness of the distribution while stretching out the lower end. Also, it makes the within-group variability more similar across groups, which in turn makes the homoscedasticity assumption by the following modeling acceptable. The second strategy is to use min-max scaling to normalize each feature’s measurements (**Eq. 2**). For each feature’s time-series of subject *i*, the minimum value of that feature gets transformed into a 0, the maximum value gets transformed into a 1, and every other value gets transformed into a decimal between 0 and 1. This normalization step is crucial to be able to emphasize the time-series pattern rather than its amplitude, which implicitly corrects for the differing baseline measurements between subjects. However, min-max normalization is not robust in handling outlier’s measurement within any time-series. Hence, outliers need to be removed as a preprocessing step prior to performing min-max normalization.

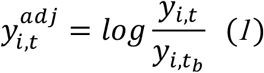

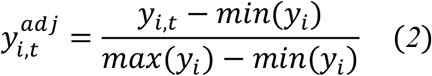

### 2.2 Fitting Gaussian smoothing spline regression model

For each omic feature *f = 1*, …, *F* from the candidate list, the data under consideration are the random variables 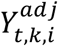 or their observations 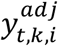 of level or mapped reads of the *i*th subject of group *k* to the feature *f* at timepoint *t*, where *t=1*, …, *T*, k=1,2, and subject *i=1*, …, *n*^*k*^. The random variable 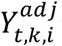 is assumed to follow 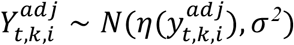, where 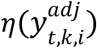 is the cubic spline function to be estimated from the data. We seek the estimation of model parameters by solving the penalized likelihood function in (**Eq.3**) using a piecewise cubic polynomial minimizer. In the objective function, ℒ = *logL*(*η*|*Y*) encourages the goodness of fit, *J*(*η*)is a roughness penalty that is added to the minus log-likelihood to quantify the smoothness of *η*, which is essentially the inner product in a reproducing kernel Hilbert space ^21^. The *λ* in (**Eq.3**) controls the trade-off between the goodness of fit and the smoothness of the spline and can be determined using cross-validation ^21^. For each feature, we solve (**Eq.3**) for each one of the two tested groups, which leads to two smoothing splines, one for each group.

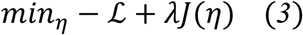

Once we have the two smoothing splines, one that fits each group’s longitudinal samples, we then calculate the test statistic for each of the *T-1* time intervals, where *T* is the number of time intervals that span the study period. We developed studentized test statistics that quantify differences between the two splines for each time interval. The formula represents the area between the two splines for each time interval *(t, t+1)* as shown in (**Eq.4**). where 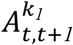 and 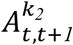 denote the area under the spline curve from time *t* to *t+1* for group 1 and group 2, respectively, *t=1*,…, *T-1*, and *SE* represents the standard error. Usually, the predicted time intervals are equidistant, as shown in **Figure 1**. Therefore, (**Eq.4**) can be rewritten in-terms of the spline function 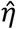 as shown in (**Eq.5**). Under the null hypothesis of no difference between the groups at the specific window, we expect the test statistics to take values near 0, with variance estimated using a permutation procedure described next.

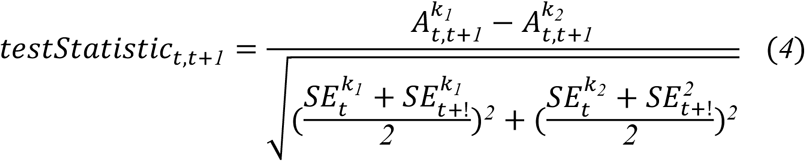

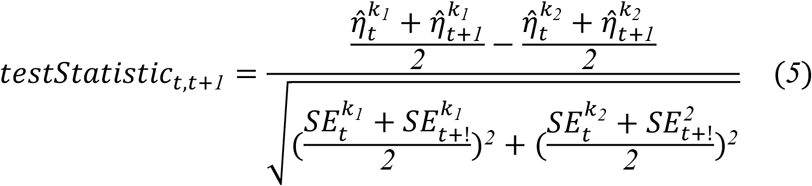

Under the null hypothesis of no difference between the groups at the specific window, we expect the test statistics to take values near 0, with variance estimated using a permutation procedure described next.

### 2.3 Inference of significant time intervals via permutation procedure

We perform a permutation procedure by permuting the sample group labels *k*. The permutation is done *B* times, and after each permutation, we calculate the 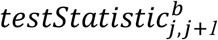 for the null hypothesis for each time interval. Since all longitudinal samples from the same participant have the same group label after each permutation, the auto-correlation correlation is preserved across subjects, and hence, type 1 error remains the same throughout the permutation procedure. Subsequently, the *pvalue* of each interval of the tested feature *f* is calculated using (**Eq.6**) when *testStatistic*_*t,t*+*1*_is positive and (**Eq.7**) when it is negative, where *T-1* denotes the number of time intervals, and *I(*.) is an indicator function. The pvalue is adjusted for multiple testing using Benjamini-Hochberg to control for the false discovery rate. For each feature *f*, significant time intervals are those with *pvalue*_*t,t*+*1*_ < *α*, where *α* is the significance level.

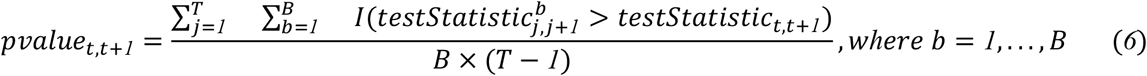

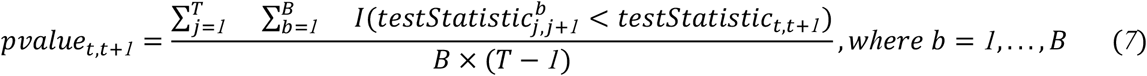

### 2.4 Global testing approach to preselect features candidates

Omics experiments usually yield profiles consisting of the expression, intensity, or abundance of thousands of molecules. The vast majority of molecules may not be temporally associated with the study groups. Hence, it is more practical to initially perform global testing on these features to select the candidate features that would be subsequently tested for time interval significance. This is a practical solution due to the computational cost of the non-parametric time interval analysis that requires a permutation procedure. Our strategy for global testing is to utilize *Edge* method ^1^ to select candidate features that show significant temporal differences between study groups. *Edge* identifies features with temporal differences during the study period without identifying when exactly the difference happens, which is the main goal of our proposed method. The *Edge* method identifies features with temporal differences by modeling each feature (**Eq.8**), where *y*_*ijkm*_ is the relative expression level of feature *i* on individual *j*, from group *m*, at the *k*th time point. The population average time curve for gene *i* of group *m* is *μ*_*im*_(*t*). Individuals deviate from the population average time curve by *γ*_*ijm*_(*t*). The measurement error and remaining sources of random variation are modeled by *ϵ*_*ijm*_(*t*). The population average time variable *μ*_*im*_(*t*) is expanded into a suitable spline basis (**Eq.9**).

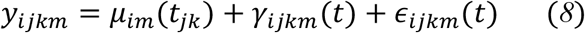

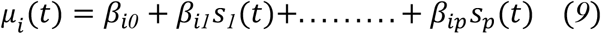

## 3. Results

### 3.1 Performance evaluation using simulated data

To measure the performance of our proposed method in identifying the significant time intervals of omics features, we simulated datasets that mimic all variations in human sample collection, such as nonuniform time gap between samples, subjects have a different number of samples over the study period, a different baseline for each subject, subjects drop out late in the study, and missing data. We used *SimStudy* ^22^ to simulate 5 datasets with different patterns across the study course, as shown in **Figure 2**. We simulated 1000 features from each pattern. The time 0 indicates the start of the study or the start of the perturbation. In our simulations, we assumed that the covariance structure between consecutive timepoints follows first-order autoregression AR(1) with a correlation coefficient ***ρ*** = 0.4. Datasets were simulated for 10 individuals from each group. Then, to mimic variability in sample collections, we sampled data points with a variable number of subjects and a variable number of samples per subject. The generated longitudinal data has a varying number of timepoints as well as varying time intervals between each measurement period. We assumed that the number of timepoints per subject follows a truncated Poisson distribution with *λ* = *20*.

**Figure 2:**
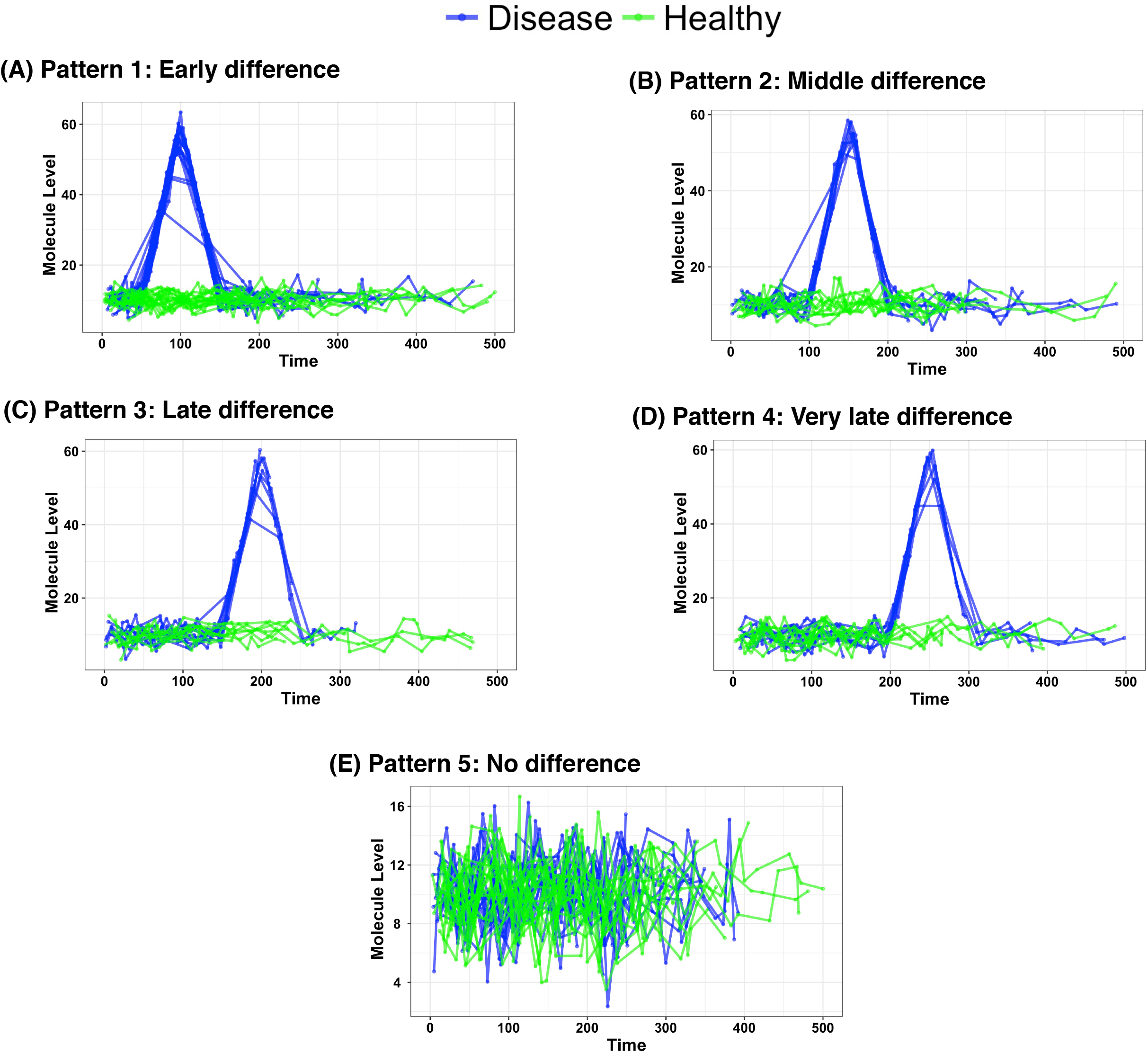
Examples of simulated features from the 5 patterns we have in this study. The first pattern indicates that the change between the two groups happened 50 days from the start of the perturbation and lasts till 150 days, pattern 2 shows differences between 100-200, pattern 3 shows differences between 150-250, pattern 4 shows differences between 200-300, pattern 5 has no change at all between the two groups and act as a negative control.

Simulated omic features of the first pattern (**Figure 2A**) were simulated with mean ***μ***(t), which follows (**Eq.10**), where ℕ denotes normal distribution, and t=0, …,500. The first pattern indicates that the change between the two groups happens 50 days from the start of the perturbation and lasts till 150 days, pattern 2 (**Figure 2B**) shows differences between 100-200, pattern 3 (**Figure 2C**) shows differences between 150-250, pattern 4 (**Figure 2D**) shows differences between 200-300, pattern 5 (**Figure 2E**) does not have change at all between the two groups and act as a negative control. The purpose of simulating these various patterns is to benchmark the proposed method performance while there are fewer samples at the period that have differences between groups since subjects dropping out of most of the longitudinal studies is directly proportional to the time.

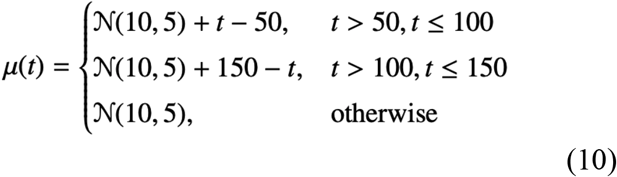

We evaluated the performance of the *OmicsLonDA* in identifying significant time intervals from each one of the 5 patterns described above. In our analysis, we used B=1000 permutations to construct the empirical distribution, a significance level of *α* = *0*.*05*, and adjusted for multiple testing using the Benjamini-Hochberg (BH) method. We tested 500 intervals for each feature (T=1, …., 501). 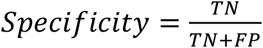 and 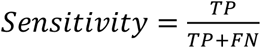,for each pattern, were measured for each feature independently, where TP is the number of significant time intervals that were correctly identified by the method, FN is the number of significant time intervals that were missed by the method, TN is the number of insignificant time intervals that were identified as insignificant, and FN is the number of significant time intervals that were identified as insignificant. Then, average specificity and sensitivity were measured among the 1000 features for each pattern. We benchmarked two variants of *OmicsLonDA* based on the model they use to fit the longitudinal data for each group; (a) OmicsLonDA with smoothing spline ANOVA (*OmicsLonDA_SSANOVA*), and (b) *OmicsLonDA* with gaussian additive mixed models (*OmicsLonDA_GAMM*). GAMM allows fitting smoothing terms to model time-series data, and it uses subject ID as a random effect. No covariates were added in this simulation study. We also benchmarked *OmicsLonDA* against *MetaLonDA*. There are three key differences between *OmicsLonDA* and *MetaLonDA*: (1) *MetaLonDA* does not correct for personal baseline, (2) *MetaLonDA* uses negative binomial smoothing spline when used with microbiome data and LOESS regression otherwise, (3) *MetaLonDA* uses a different formula for testStatistic that only include the area between the curves of the two groups without adjusting of the standard error in their estimation.

**Figure 3A** demonstrates the high level of specificity of *OmicsLonDA* (>0.99) among all 5 tested patterns, with *OmicsLonDA_GAMM* has slightly more specificity over the first 4 patterns, and *OmicsLonDA_SSANOVA* has slightly more specificity in Pattern-5. On the other hand, *MetaLonDA’s* specificity is ∼0.80 among all first 4 patterns, and 0.97 in pattern-5. **Figure 3B** shows the sensitivity of all benchmarked methods for all patterns, except pattern-5. This is because all features in pattern-5 were simulated to not have any significant differences in the time intervals between the two compared groups. *MetaLonDA* has the highest sensitivity (∼0.98) among the compared methods. This high sensitivity can also be seen as a trade-off with the low specificity of *MetaLoNDA* shown in **Fig 3A**. The sensitivity for all methods from pattern-1 to pattern-4. This decrease in sensitivity is expected due to the fact that as the time intervals that are significantly different between the two groups shift to the right (later in the study course, which was implemented in our simulations), there are more participants dropping out of the study, and hence there is lower power of each method to detect the significantly differential time intervals. *OmicsLonDA_SSANOVA* maintains reasonably high sensitivity across all patterns (pattern-1: 0.98, pattern-1: 0.92, pattern-3: 0.90, and pattern-4: 0.87). *OmicsLonDA_GAMM* has a similar sensitivity pattern to *OmicsLonDA_SSANOVA*, but surprisingly, the sensitivity drops significantly at pattern-4 (0.72). These results demonstrate that *OmicsLonDA_SSANOVA* is a better choice than *OmicsLonDA_GAMM* when there are few samples covering the tested time intervals.

**Figure 3:**
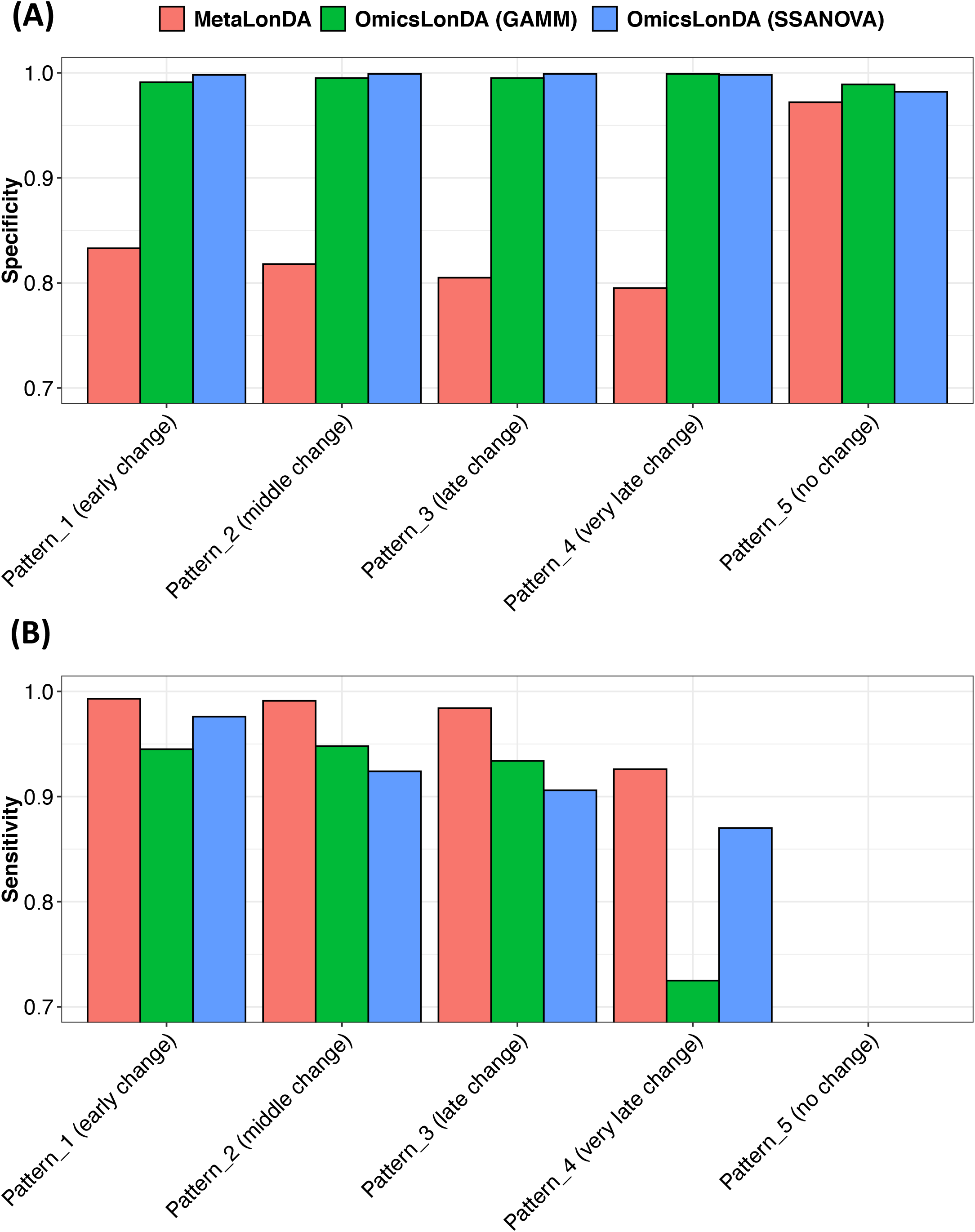
Performance evaluation of identifying significant time intervals from simulated features.

Additionally, we evaluated the two implemented personal baseline adjusting methods (log-ratio, and min-max normalization) on *OmicsLonDA* performance. Table 1 shows the sensitivity and specificity of *OmicsLonDA* when ran on the 5 simulated patterns after each of the baseline adjusting methods. While log-ratio and min-max baseline adjusting methods have yielded a similar effect on *OmicsLonDA* specificity, min-max yields higher sensitivity.

**Table 1:** Evaluation of adjusting subject’s profile.

|  | *OmicsLonDA* (log-ratio) |  | *OmicsLonDA* (min-max) |  |
| --- | --- | --- | --- | --- |
|  | Sensitivity | Specificity | Sensitivity | Specificity |
| Pattern_1 (early change) | 0.97 | 0.96 | 0.97 | 0.99 |
| Pattern_2 (middle change) | 0.93 | 0.96 | 0.93 | 0.99 |
| Pattern_3 (late change) | 0.88 | 0.98 | 0.9 | 0.99 |
| Pattern_4 (very late change) | 0.81 | 0.99 | 0.88 | 0.99 |
| Pattern_5 (no change) | - | 0.98 | - | 0.99 |

### 3.2 Performance evaluation of a global testing approach to pre-select features candidates

We evaluated the ability of the global testing procedure in acting as a selection criterion and providing candidates that have significant time intervals between the two interest groups. We benchmarked the previously described *Edge* method to a linear mixed effect model. We use the basic linear mixed effect model in global testing, 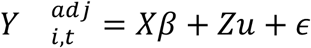, where 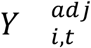 is an Nx1 column vector representing the level of *N* omic features; X is an Nxp matrix of p covariates; *β*is a px1 column vector of the fixed-effect regression coefficients; Z is the Nxq design matrix for the q random effects (subjects random effect in our case); u is a qx1 vector of the random effects, and ε is an Nx1 column vector of the residuals. **Table 2** summarizes the performance of the global testing procedure in terms of 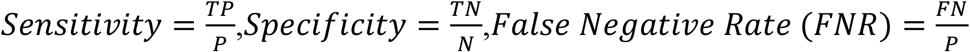, and 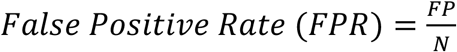. Based on our simulation, for each simulated feature from pattern 1-4, it has 100 positive time intervals. On the other hand, features simulated from pattern 5, have 0 positive time intervals since no difference is expected between the two tested groups. **Table 2** demonstrates that, for most patterns except pattern 3, it can be used as a selection criterion for candidate features to be tested for time interval analysis. As shown, although the global testing finds the majority of the features that have significant time intervals, except for pattern 3, it does not find them all. This is another strength of the time interval analysis because global testing in longitudinal studies is a compelling approach only when the features demonstrate the difference between the study groups during all the study courses and not only a portion of it. This is a trade-off to be taken into account while weighing computational time over sensitivity.

**Table 2:** Evaluation of global testing in selecting candidate features for subsequent time interval analysis.

|  | LMM |  |  | Edge |  |  |
| --- | --- | --- | --- | --- | --- | --- |
|  | # Candidates | FNR | FPR | # Candidates | FNR | FPR |
| Pattern_1 (early change) | 991 | 0.009 |  | 994 | 0.006 |  |
| Pattern_2 (middle change) | 898 | 0.102 |  | 990 | 0.010 |  |
| Pattern_3 (late change) | 994 | 0.006 |  | 1000 | 0.000 |  |
| Pattern_4 (very late change) | 997 | 0.003 |  | 989 | 0.011 |  |
| Pattern_5 (no change) | 52 |  | 0.052 | 40 |  | 0.04 |

### 3.3 Time and memory evaluation

The running time of *OmicsLonDA* depends primarily on the number of permutations used to construct the empirical distribution for each feature. In our analysis of the simulated data with 1000 permutations, for each feature, *OmicsLonDA* analysis without the global testing took on average 443 min and 47 seconds. Global testing using Edge ^1^ using 1000 bootstraps took on average 64 seconds for each feature. The evaluation was conducted on a MAC machine with a 2.5 GHz Intel Core i7 processor and 16 GB 1600 MHz RAM. Although *OmicsLonDA* supports parallel computing, we used 1 thread in our evaluation.

### 3.4 Application of *OmicsLonDA* on real-world datasets

#### 3.4.1 iPOP infection multi-omics cohort

As an application for our proposed method, we used the integrative Personal Omics Profiling (iPOP) cohort, a longitudinal cohort that aims to characterize the complex host-microbial interactions in Type 2 diabetes mellitus (T2DM) ^20^. The iPOP cohort was established to better understand T2DM at its earliest stages, where healthy or prediabetic individuals are sampled over ∼4 years in a deep multi-omics profiling of transcriptomes, metabolomes, proteomes, and cytokines, as well as gut and nasal microbiome. In a total of 1091 visits, 105 participants (25-75 years old, BMI of 19-41 kg/m^2^, 55 females and 50 males) were profiled during healthy periods and extensively during periods of respiratory viral infection (RVI), immunization, and other situations that perturb human host-microbial physiology.

We leveraged the power of the longitudinal multi-omics nature of the iPOP study to reveal sexual dimorphism at the molecular level following RVI episodes. Sex is considered to be an important epidemiological factor that can determine the risk for some diseases. However, the sex dependentt responses to respiratory viral infections are not well explored, especially in a multi-omics and microbiological fashion. Most of the previous studies were based on epidemiological strategy and reported the prevalence of RVI in different sex ^23–26^. In this work, we utilized *OmicsLonDA* to identify longitudinal transcriptomic, metabolomic, cytokines, and microbial changes between females and males following RVI. In the context of this work, we included 25 (12 male and 13 female) participants who were followed before and after RVI (44 episodes of RVI in a total of 180 RVI visits (**Figure 4**). We selected episodes that have at least 3 samples during the first 39 days after RVI. We first adjusted each feature using min-max normalization (**Eq.2**). Each feature (gene, protein, metabolite, cytokine, or microbe) was tested independently. Time interval inference is based on an empirical distribution that is built for all intervals of the same feature as described previously. We tested 38 intervals (between day 1 and day 39). For genes, out of 10346 available gene expressions, we tested only 331 that show temporal significance using global testing criteria described in the method section (qvalue<0.1). For proteins, metabolites, cytokines, and microbes, we tested all quantified features from the iPOP cohort. In our analysis, we used B=1000 permutations to construct the empirical distribution, significance level *α* = *0*.*05*, and adjusted for multiple testing using Benjamini-Hochberg (BH) method.

**Figure 4:**
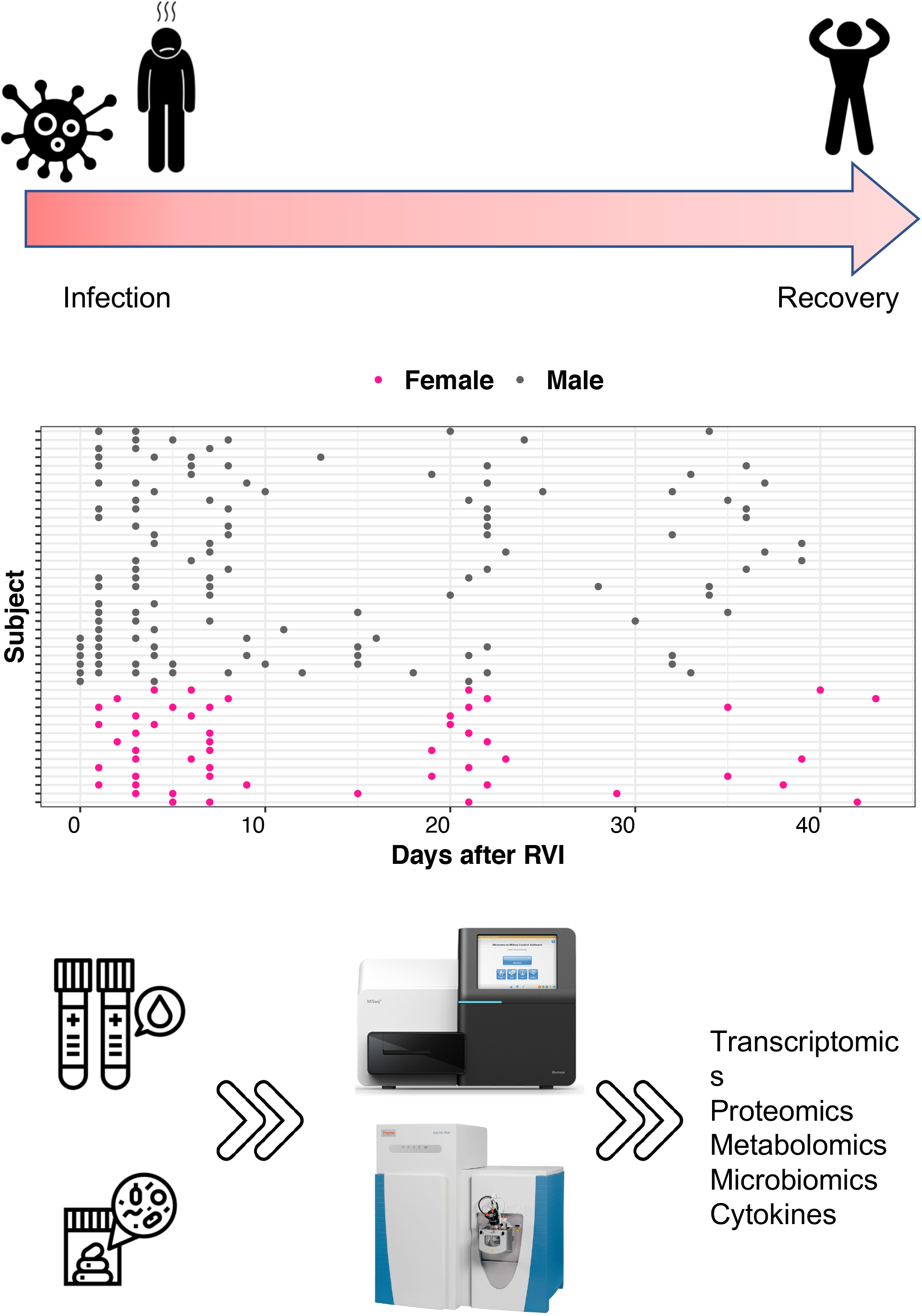
Study design of the iPOP infection cohort. Time points distributions of 44 infection episodes whose corresponding subject has at least three timepoints within 40 days following and infection incidence. Total of 180 samples from 25 subjects (12 male and 13 female). Timeline annotation of RVI episodes, where day 0 is the first day of infection.

In total, 104 features (36 genes, 29 proteins, 35 metabolites, 3 cytokines, 1 microbe) exhibit temporal differences between males and females following RVI. **Figure 5** shows a timeline summary of omic features that show the difference between males and females after RVI episodes (**Table S1**). The results reveal that females were more responsive to RVI with 58 omic features being overexpressed, while 44 features were over-expressed in males, and 2 genes (MFSD7 and SCN5A) flipped their overexpression trajectory during the course of the infection episode. Females have stronger antibody responses IGLV3-19 (LV319) and IGHV3-53 (HV353). Females have a stronger adaptive response early than males, while males have more innate responses than females with increased complement proteins and increased red blood cells (HBA1, HBB, HBD). Interestingly, males have over-expression of Vitamin D3 (Dihydroxyvitamin) during the early infection period (day 1 to day 21). Females have higher leptin during the whole course of infection (1-39 days).

**Figure 5:**
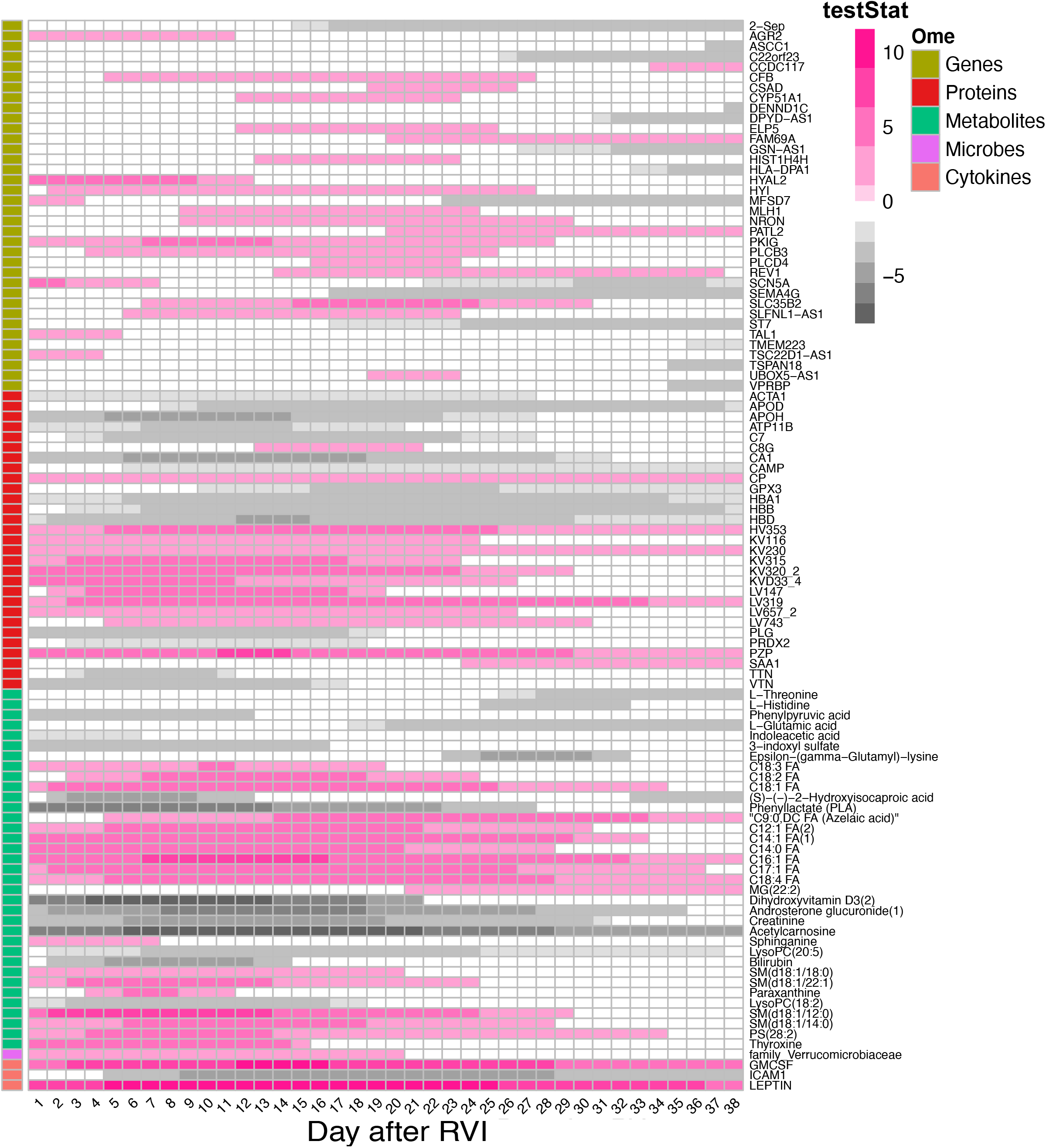
Significant time intervals of features that show differences between males and females following RVI. Each row represents a feature. Pink shaded cells indicate the corresponding feature is over-expressed in the female group, while the gray lines indicate the corresponding metabolite is over-expressed in the male group.

#### 3.4.2 Preeclampsia lipidomics cohort

We applied *OmicsLonDA* on a longitudinal lipidomics study on preeclampsia ^27^ as a case study to demonstrate the value of *OmicsLonDA* for the identification of time intervals that lipids are significantly different between pregnancy with and without preeclampsia. Preeclampsia is a serious pregnancy complication affecting 5-10% of pregnant women, accounting for approximately 40% of fetal deaths worldwide. It not only harms maternal health but also inhibits fetal growth and causes babies to be born with immature development. Therefore, detection of preeclampsia biomarkers at early gestational age and identification of the time intervals with dramatic lipid changes in preeclampsia is crucial for preeclampsia early diagnosis and treatment.

In this longitudinal prospective study, the cohort was previously described ^27^ with 27 and 20 women with and without preeclampsia, respectively. The plasma samples were collected from each subject at two or three-time points during pregnancy. The gestational age distribution of the preeclampsia and control groups in each trimester is shown in **Figure 6A**. For each plasma sample, we conducted target lipidomics analysis by applying the Lipidyzer platform for 750 lipid species composed of 13 diverse lipid classes (**Figure 6B**).

**Figure 6:**
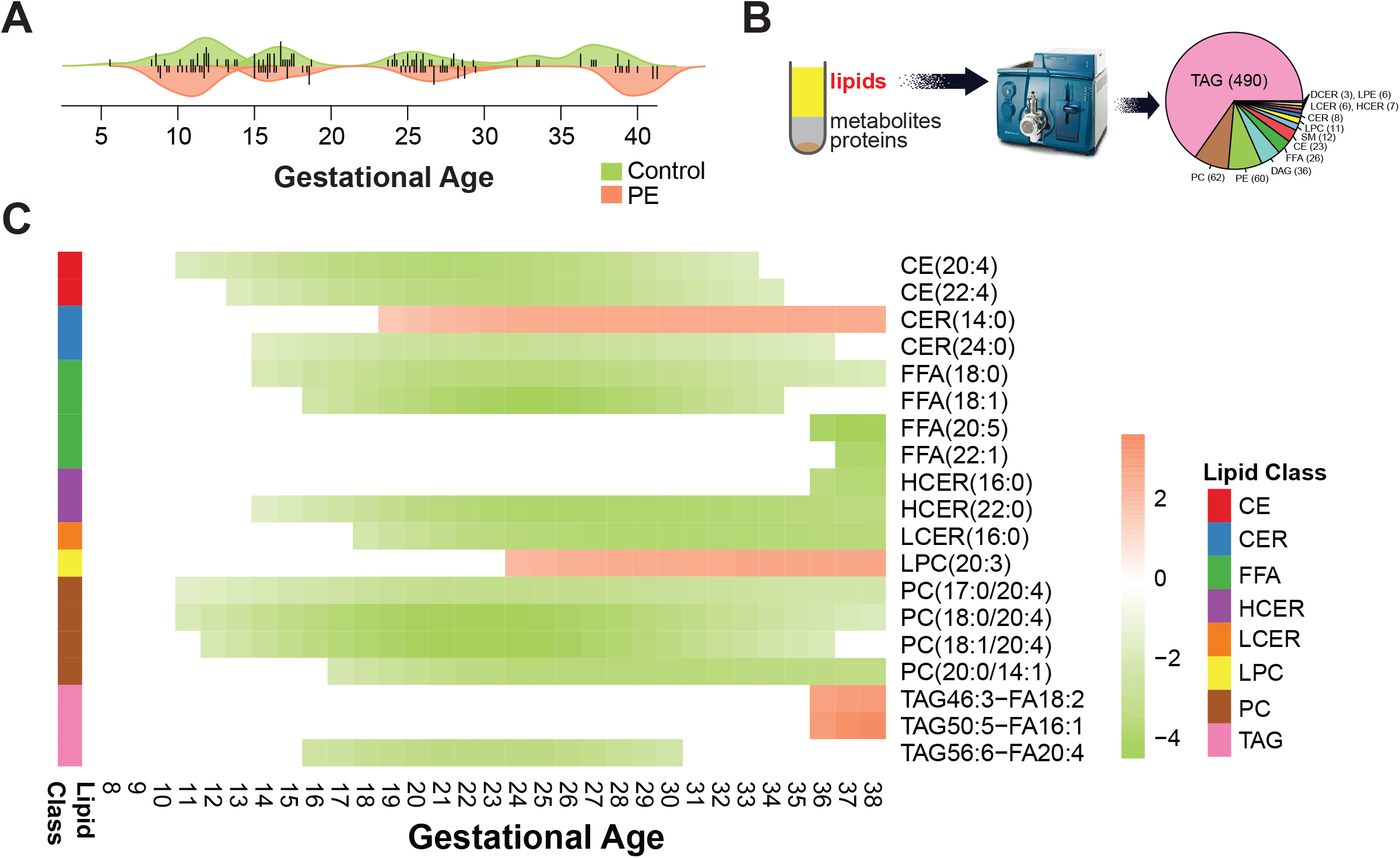
A case study of the longitudinal lipidomics data on the preeclampsia cohort. (A) The gestational age distribution of the collected plasma samples from the control and preeclampsia (PE) groups in each trimester in this study. (B) Lipids from the plasma samples were extracted and measured by the target lipidomics profiling platform Lipidyzer for 750 different lipids. (C) The 19 lipid species exhibit significantly different profiles between the control and preeclampsia groups by applying *OmicsLonDA*.

Following *OmicsLonDA* analysis workflow, we first adjusted the levels of each lipid at later time-points using min-max normalization method (Eq.2) and normalized them to baseline. The *OmicsLonDA* test was conducted for each lipid independently. We set one week as one time interval unit and tested on 30 time intervals (week 8-38). In our analysis, we used 1,000 times permutations to construct the empirical distribution for each lipid. All the results were adjusted for multiple testing using Benjamini-Hochberg (BH) method with a significance level *α* = *0*.*05*.

We identified 19 lipid species, accounting for eight lipid classes, with significant temporal differences between preeclampsia and control groups during pregnancy. **Figure 6C** demonstrates the time intervals of these 19 lipids that have significantly different profiles between preeclampsia and control groups (**Table S2**). Interestingly, we found that most of the significant lipids belonging to the same lipid classes exhibit the same changing trends, indicating homogeneity of chemical properties and potential biological roles of lipid species from the same classes. There are two exceptions: one is CER (14:0) and CER (24:0), which may be due to the different lengths of fatty acid chains in these two ceramides. Another interesting exception is TAG 46:3 (FA 18:2), TAG 50:5 (FA16:1), and TAG 56:6 (FA20:4), showing the higher levels of the first two triglycerides in the preeclampsia group at later gestational age compared to the control group. In contrast, TAG 56:6 (FA20:4) displays the increased abundance at early pregnancy in control subjects. Further experiments are needed to investigate the explicit reasons.

## 4. Discussion

In this work, we have developed a statistical method that provides robust identification of time intervals where omics features are significantly different between groups in longitudinal multiomics. The method is able to simultaneously identify time intervals and differential signatures by analyzing each feature separately, but across all patients. The proposed method is based on a semi-parametric approach, where using smoothing splines to model longitudinal data and infer significant time intervals of omics features based on an empirical distribution constructed through the permutation procedure. A critical need in longitudinal omics is for robust frameworks that incorporate the time dimension in statistical significance analysis. Our method, evaluated through extensive simulations (5 patterns). The performance evaluation demonstrated that *OmicslonDA* has achieved a correctly calibrated type-I error rate and is robust to data collection inconsistencies that commonly occur in longitudinal human studies. Moreover, the sensitivity is high in pattern 1 and then declines slightly through pattern 4. This decrease in sensitivity can be explained due to the decreasing number of samples collected towards the end of the study (i.e., patient dropout). We further applied *OmicsLonDA* on two real-world datasets: (1) the iPOP longitudinal omics study for investigating sexual dimorphism on molecular response following respiratory viral infection, (2) preeclampsia cohort to identify time intervals that lipids are significantly different between pregnancy with and without preeclampsia. Recently, OmicsLonDA has been utilized to identify the seasonal time intervals of differentially abundant/expressed omics features between insulin-resistant and insulin-sensitive individuals ^28^.

Sex differences in response to infection are known ^29^. For both viral and bacterial infections, males are more susceptible than females, while females produce a more vigorous inflammatory response ^29^. Sexual dimorphism in infection response likely arises from differences in hormone status, with both testosterone and estrogen shown to modulate infection and inflammatory processes ^30^. Our study further adds evidence that sexual dimorphism may contribute to stages of inflammatory responses ^31^, with females having a stronger adaptive response early but less innate responses than male. Our analysis is the first to our knowledge that revealed a multitude of sex differences in RSV infection response with frequent temporal sampling and delineation of the dynamic infection response for each sex.

Preeclampsia is a potentially life-threatening complication during pregnancy identified by increased blood pressure and proteinuria. It is one of the leading causes of maternal and perinatal mortality and morbidity ^32^. Substantial efforts have been made to detect molecular changes of preeclampsia during pregnancy at gene, protein and metabolite levels ^33–35^. Nowadays, lipids are growingly recognized as key players involved in pathophysiology of preeclampsia ^36,37^. For instance, arachidonic acid and its downstream products were reported to be significantly changed in preeclampsia ^38^. Oxidized lipid species were also selected as biomarkers of preeclampsia which are related to increased reactive oxygen species (ROS) ^39^. However, most of these studies mainly focused on single timepoint instead of monitoring dynamic molecular changes with multiple timepoints during pregnancy. Herein, in this study we applied the developed *OmicsLonDA* on a longitudinal lipidomics dataset to compare lipid levels in women with and without preeclampsia. We successfully identified 19 distinct lipid species that are significantly different in preeclampsia pregnancy compared to normal pregnancy with different time intervals. Interestingly, most of the significant lipids that belong to the same lipid classes exhibit the same changing trends, indicating potentially similar biological functions these lipids may exert in preeclampsia progression. Importantly, by *OmicsLonDA*, we detected several lipids that harboured significantly different time intervals at early pregnancy phase (i.e., the first trimester) such as CE (20:4), PC (17:0/20:4), PC (18:0/20:4) and PC (18:1/20:4), which may serve as clinically meaningful biomarkers for preeclampsia early diagnosis. Intriguingly, all of these four lipid biomarkers share the same fatty acid chain arachidonic acid (fatty acid 20:4). Arachidonic acid is a polyunsaturated fatty acid containing 20 carbons and four double bonds with a final double bond in the ω−6 position. It is well-documented that arachidonic acid and its products eicosanoids play important roles in inflammatory processes ^40^. They have been reported as biomarkers of preeclampsia previously ^38^. Our results not only support the previous findings, but also revealed more potentially involved lipid biomarkers by leveraging the advantages of the longitudinal data as well as the merit of *OmicLonDA*.

*OmicsLonDA* elucidates not only differentially regulated molecules but indicates the temporal window over which the differential regulation occurs to provide a nuanced and detailed understanding of biological dynamics. In the future, we plan to utilize the identified multi-omics features and their significant time intervals through non-parametric Bayesian dynamic networks to infer causality of phenotypes based on a phased correlation between features’ time intervals and phenotype onset. Another avenue for improving the proposed method is to incorporate autocorrelation between longitudinal samples into the model fitting. Also, in the proposed method, time intervals to be tested are a user-defined parameter. In the future, we plan to develop a learning method that selects non-trivial intervals that span several timepoints. *OmicsLonDA* is publicly available on the Bioconductor repository (https://bioconductor.org/packages/OmicsLonDA).

## Supplementary Material

**Table S1:** *OmicsLonDA* results of iPOP infection cohort. Rows represent different omics features that showed temporal significant differences between males and females after RVI, columns represent time intervals in days, and each cell represents the testStat for significant time intervals and zero otherwise.

**Table S2:** *OmicsLonDA* results of preeclampsia. Rows represent different lipid features that showed temporal significant differences between males and females after RVI, columns represent time intervals in days, and each cell represents the testStat for significant time intervals and zero otherwise.

## Acknowledgments

We thank Robert Tibshirani from Stanford University for his valuable suggestions and critical comments throughout the development of *OmicsLonDA*. We would like to thank iPOP participants and numerous colleagues from Stanford University who contributed to the sample collection, preparation, and processing of the iPOP cohort.

## Data and Code Availability

*OmicsLonDA* is publicly available on the Bioconductor repository (https://bioconductor.org/packages/OmicsLonDA). The development source code is located on Github (https://github.com/aametwally/OmicsLonDA). iPOP data is publicly available at the human microbiome project portal (https://portal.hmpdacc.org/).

